# Chordin-mediated BMP shuttling patterns the secondary body axis in a cnidarian

**DOI:** 10.1101/2024.05.27.596067

**Authors:** David Mörsdorf, Maria Mandela Prünster, Grigory Genikhovich

## Abstract

BMP signaling patterns secondary body axes throughout Bilateria and, strikingly, in the bilaterally symmetric corals and sea anemones – members of the bilaterian sister clade Cnidaria. It has been suggested that the secondary, “directive” axis in the sea anemone *Nematostella vectensis* requires Chordin-mediated “shuttling” of BMP ligands, like in *Drosophila* or frog, however, an alternative “local inhibition” model is also possible. To choose between these two options, we generated localized Chordin sources in the Chordin morphant background and showed that in the presence of BMP ligands in *Nematostella*, mobile Chordin is necessary and sufficient to establish a peak of BMP signaling at the side of the embryo opposing the Chordin source. In contrast, membrane-tethered Chordin-CD2 promotes weak BMP signaling within the Chordin-CD2 source. These results provide the first mechanistic evidence for BMP shuttling in a cnidarian and suggest that BMP shuttling may have been functional in the cnidarian-bilaterian ancestor.

## Introduction

Bone Morphogenetic Protein (BMP) signaling acts in secondary body axis patterning across Bilateria, and its functions as morphogen have been studied in diverse animal species (Bier and De Robertis, 2015; Mörsdorf et al., 2024). The mechanisms of the BMP-dependent axial patterning are remarkably similar between arthropods and vertebrates, indicative of the shared origin of the secondary, dorsoventral axis in protostome and deuterostome Bilateria – a notion strengthened once broader phylogenetic sampling became available (Ben-Zvi et al., 2008; Dale et al., 1992; Ferguson and Anderson, 1992; Holley et al., 1995; Irish and Gelbart, 1987; Mörsdorf et al., 2024). Intriguingly, the same mechanisms appear to regulate the secondary axis patterning in the bilaterally symmetric cnidarian *Nematostella vectensis*, indicating that a BMP-dependent secondary body axis may have evolved prior to the evolutionary split of Cnidaria and Bilateria (Matus et al., 2006; Saina et al., 2009; reviewed in Bier and De Robertis, 2015; Genikhovich and Technau, 2017). However, an alternative scenario, that BMP-mediated secondary axis evolved convergently in Bilateria and bilaterally symmetric Cnidaria, is also possible (Mörsdorf et al., 2024).

BMPs are secreted signaling proteins of the Transforming Growth Factor-β superfamily frequently acting as heterodimers (Bauer et al., 2023; Little and Mullins, 2009; Tajer et al., 2021). Signaling through the BMP receptor complex (Fig. 1A) results in phosphorylation and nuclear accumulation of the transcriptional effector SMAD1/5, which regulates the expression of many crucial developmental transcription factors and signaling pathway components (Deignan et al., 2016; Greenfeld et al., 2021; Knabl et al., 2024; Rogers et al., 2020; Stevens et al., 2017; reviews by Akiyama et al., 2024; Hill, 2016). BMP signaling is tightly controlled by a plethora of intracellular (Knabl et al., 2024; Miller et al., 2019) and extracellular regulators (Chang et al., 2001; Dal-Pra et al., 2006; Hsu et al., 1998; Piccolo et al., 1997; Ross et al., 2001; Sasai et al., 1994; Scott et al., 2001; Zimmerman et al., 1996). Chordin (= Short gastrulation in insects) is, arguably, the most famous extracellular regulator of BMP signaling. Like many other BMP antagonists, Chordin binds BMP ligands, blocks the interaction with their receptor and thereby inhibits BMP signaling (Piccolo et al., 1996). However, Chordin can also have pro-BMP effects and promotes long-range activation of BMP signaling in *Drosophila*, *Xenopus*, sea urchins and also in *Nematostella* (Ashe and Levine, 1999; Ben-Zvi et al., 2008; Genikhovich et al., 2015; Lapraz et al., 2015; Wang and Ferguson, 2005). The phylogenetic distribution of Chordin and two central BMP ligands, BMP2/4 and BMP5-8, and their importance for the secondary axis patterning across phyla suggests that during early animal evolution, these molecules may have represented the minimum requirement for the formation of the bilaterally symmetric body plan (Genikhovich and Technau, 2017; Mörsdorf et al., 2024). However, to evaluate such a possibility, we need to understand the “mode of action” of BMPs and Chordin outside Bilateria, and our model, the sea anemone *Nematostella*, allows exactly that.

**Figure 1:**
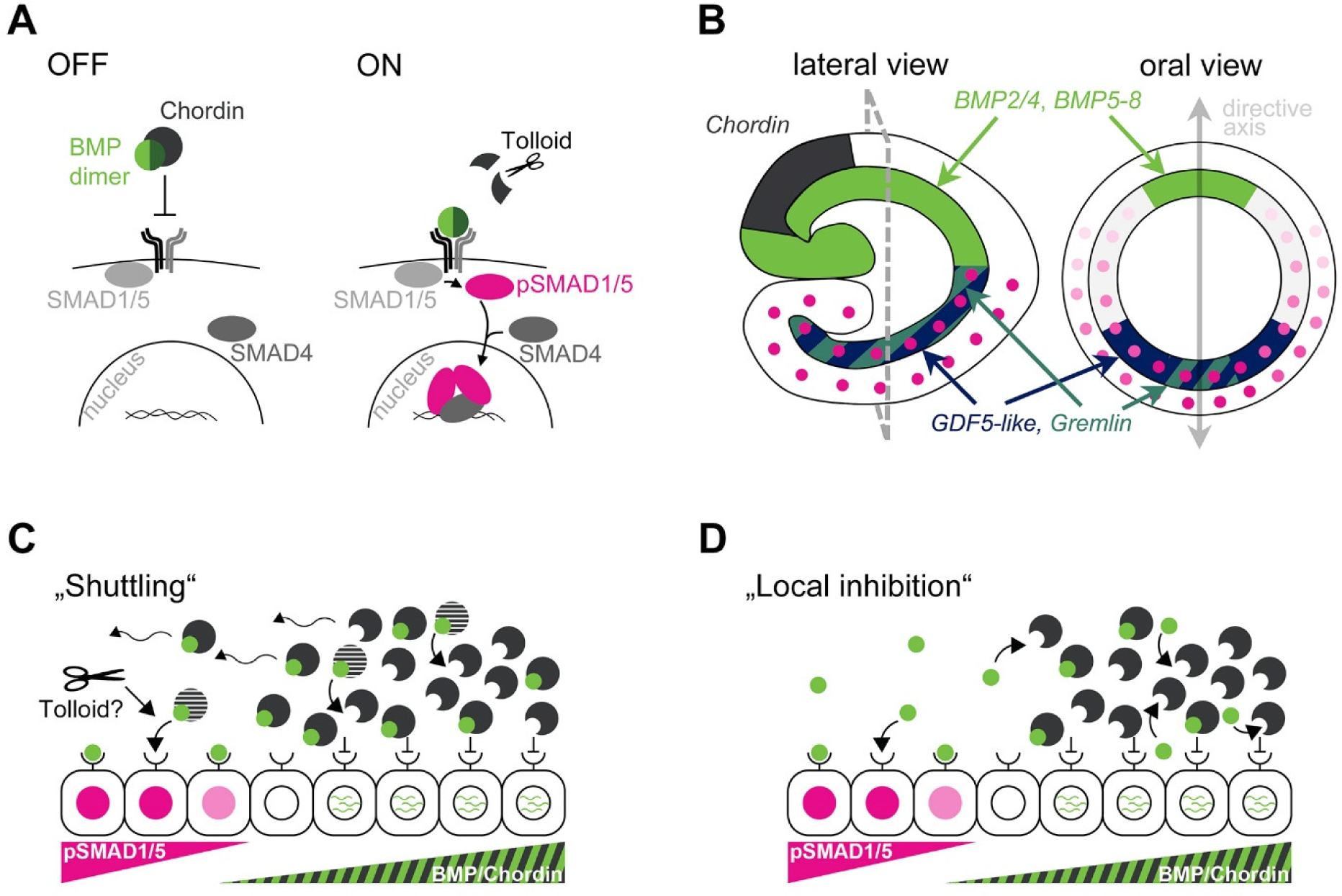
Possible modes of action of BMP signaling during axial patterning in *Nematostella*. **A)** BMP signaling pathway. BMP dimers bind the heterotetrameric receptor complex, resulting in the phosphorylation of SMAD1/5. pSMAD1/5 forms a complex with the Co-Smad SMAD4, which regulates transcription in the nucleus. Chordin binds BMPs preventing them from activating the receptor complex. Metalloproteases like Tolloid and BMP-1 cleave Chordin and release BMP ligands from the inhibitory complex in Bilateria. **B)** Expression domains of BMPs and BMP antagonists in an early *Nematostella* larva. Oral view corresponds to the optical section indicated with grey dashed line on the lateral view. Pink circles show the nuclear pSMAD1/5 gradient. **C)** The shuttling model suggests that in *Nematostella*, a mobile BMP-Chordin complex transports BMPs through the embryo. Receptor binding is inhibited in cells close to the Chordin source due to high concentrations of Chordin. On the opposite side of the directive axis, BMPs bind their receptors and activate signaling upon release from Chordin. Tolloid might be involved in the cleavage of Chordin and release of BMPs from the complex with Chordin also in *Nematostella*. **D)** In the local inhibition model, *Nematostella* Chordin acts locally to inhibit BMP signaling and promote the production of *BMP2/4* and *BMP5-8* mRNA. Chordin mobility is not required for asymmetric BMP signaling.

BMP signaling in *Nematostella* becomes detectable during early gastrula stage in a radially symmetric domain: pSMAD1/5 is detected in the nuclei around the blastopore (Knabl et al., 2024; Leclère and Rentzsch, 2014). Shortly after the onset of BMP activity, the radial symmetry of the embryo breaks, establishing the secondary, “directive” body axis with minimum BMP signaling intensity detectable on the side of *BMP2/4*, *BMP5-8* and *Chordin* expression and maximum BMP signaling on the side opposite to it (Fig. 1B; Genikhovich et al., 2015; Knabl et al., 2024; Leclère and Rentzsch, 2014). Both BMP2/4 and BMP5-8 are crucial for BMP signaling and directive axis patterning since knockdown of either ligand abolishes the BMP signaling gradient (Genikhovich et al., 2015). While this finding suggests that BMP2/4 and BMP5-8 signal as an obligate heterodimer during axial patterning in *Nematostella*, BMP2/4 and BMP5-8 are not the only BMP ligands present in the embryo at this stage. GDF5-like, a BMP ligand that is expressed on the side of strong BMP signaling (Fig. 1B), appears to steepen the pSMAD1/5 gradient (Genikhovich et al., 2015; He et al., 2023; Knabl et al., 2024). The BMP signaling gradient is stable over many (>24) hours during which it patterns the directive axis (Genikhovich et al., 2015; He et al., 2023; Knabl et al., 2024; Leclère and Rentzsch, 2014; Saina et al., 2009). Considering the short half-life of phosphorylated SMAD1/5 reported in other systems (Miller et al., 2019; Rogers et al., 2020), this indicates that long-range transport (∼100 µm) of BMP2/4 and BMP5-8 and constant receptor complex activation is necessary to maintain BMP signaling. How it exactly happens that the core BMP ligands, BMP2/4 and BMP5-8, are expressed on one side of the embryo and the peak of BMP signaling activity is on the opposite side is currently unknown.

One possible explanation involves Chordin-mediated shuttling of BMP ligands, described in the dorsoventral patterning in *Drosophila* and *Xenopus* (Ben-Zvi et al., 2008; Genikhovich et al., 2015; Mizutani et al., 2005). In this model, Chordin inhibits BMP function locally, close to the Chordin source cells, but promotes long-range BMP signaling by forming a mobile complex with the BMP dimer, which is released once Chordin is cleaved by the metalloprotease Tolloid. The probability that this BMP dimer will bind its receptors rather than another, yet uncleaved Chordin increases with the distance to the Chordin source (Fig. 1C). In *Nematostella*, the shuttling model is supported by the finding that, unlike in all bilaterian models studied thus far, depletion of Chordin results in the loss of BMP signaling (Genikhovich et al., 2015). However, given that BMP signaling indirectly represses *BMP2/4* and *BMP5-8* transcription at this developmental stage (Knabl et al., 2024), an alternative explanation that Chordin locally represses BMP signaling enabling BMP2/4 and BMP5-8 production and diffusion into the area of low or no Chordin (*i.e.* to the *GDF5-like* side of the directive axis), is also possible. This “local inhibition” model in which Chordin acts exclusively as a local repressor of BMP signaling (Fig. 1D) is similar to the situation in zebrafish, where Chordin does not show any long-range pro-BMP effects and extracellular transport of Chordin is not required (Pomreinke et al., 2017; Tuazon et al., 2020; Zinski et al., 2017). In this paper, we address the role of Chordin in the BMP-dependent axial patterning in the sea anemone *Nematostella* and test these two alternative models.

## Results

### BMPs are retained on the surface of embryonic cells

We reasoned that the most straightforward way of addressing the role of Chordin in BMP signaling was to visualize active BMPs in the presence or absence of Chordin. To that end, we set out to generate biologically active, detectable BMP2/4, express it in the endogenous domain and address its distribution. A critical step of the posttranslational processing of BMP ligands is the proteolytic cleavage of the BMP propeptides, separating the prodomain from the mature domain (Akiyama et al., 2024). Therefore, we generated a tagged BMP2/4, in which a FLAG-superfolder GFP (sfGFP) tag is fused to the N-terminus of the mature domain. This design is based on previous BMP fusion proteins (Pomreinke et al., 2017) and results in a BMP ligand that can activate BMP signaling, as shown by pSMAD1/5 western blot (Fig. 2A, Supplementary Fig. 1A). Co-injection of *BMP2/4* and *BMP5-8* mRNAs leads to a stronger phosphorylation of SMAD1/5 than does the injection of individual ligand mRNAs (Supplementary Fig. 1B,C), indicating that heterodimers consisting of BMP2/4 and BMP5-8 are indeed the relevant BMP ligands *in vivo*, consistent with their identical knockdown phenotypes (Genikhovich et al., 2015; Saina et al., 2009). Then, we generated a transgenic line in which *BMP2/4-FLAG-sfGFP* is expressed under control of the 4.5 kb DNA fragment upstream of the start codon of *Nematostella BMP2/4* (Schwaiger et al., 2014). An F0 male with germline-transmission of the transgene was crossed to a wildtype female and pSMAD1/5-positive nuclei were detected by immunofluorescence in GFP-positive F1 embryos to visualize the BMP2/4-FLAG-sfGFP protein and the BMP signaling domain simultaneously. Consistent with the previously described expression of endogenous *Nematostella bmp2/4* on the “low BMP signaling” side of the secondary body axis (Genikhovich et al., 2015), we observed a graded GFP signal with the maximum on the pSMAD1/5-negative side of the 2 days post fertilization (dpf) embryo (Fig. 2B). Due to several silent mutations, the translation of the BMP2/4-FLAG-sfGFP is not affected by the previously characterized BMP2/4 morpholino (BMP2/4MO; Genikhovich et al., 2015; Kirillova et al., 2018; Knabl et al., 2024; Saina et al., 2009), which allowed us to test the activity of the *BMP2/4-FLAG-sfGFP* transgene in the absence of endogenous BMP2/4. The transgene only partially rescued the phenotype of BMP2/4MO-injected embryos, and none of them formed primary polyps, suggesting that some necessary enhancer elements were missing in our construct. However, pSMAD1/5-positive cells could be detected on one side of the directive axis in approximately 1/3 of the transgenic BMP2/4MO-injected embryos (9/31; Fig. 2C), while all non-transgenic BMP2/4MO-injected controls remained pSMAD1/5-negative (12/12; Fig. 2C).

**Figure 2:**
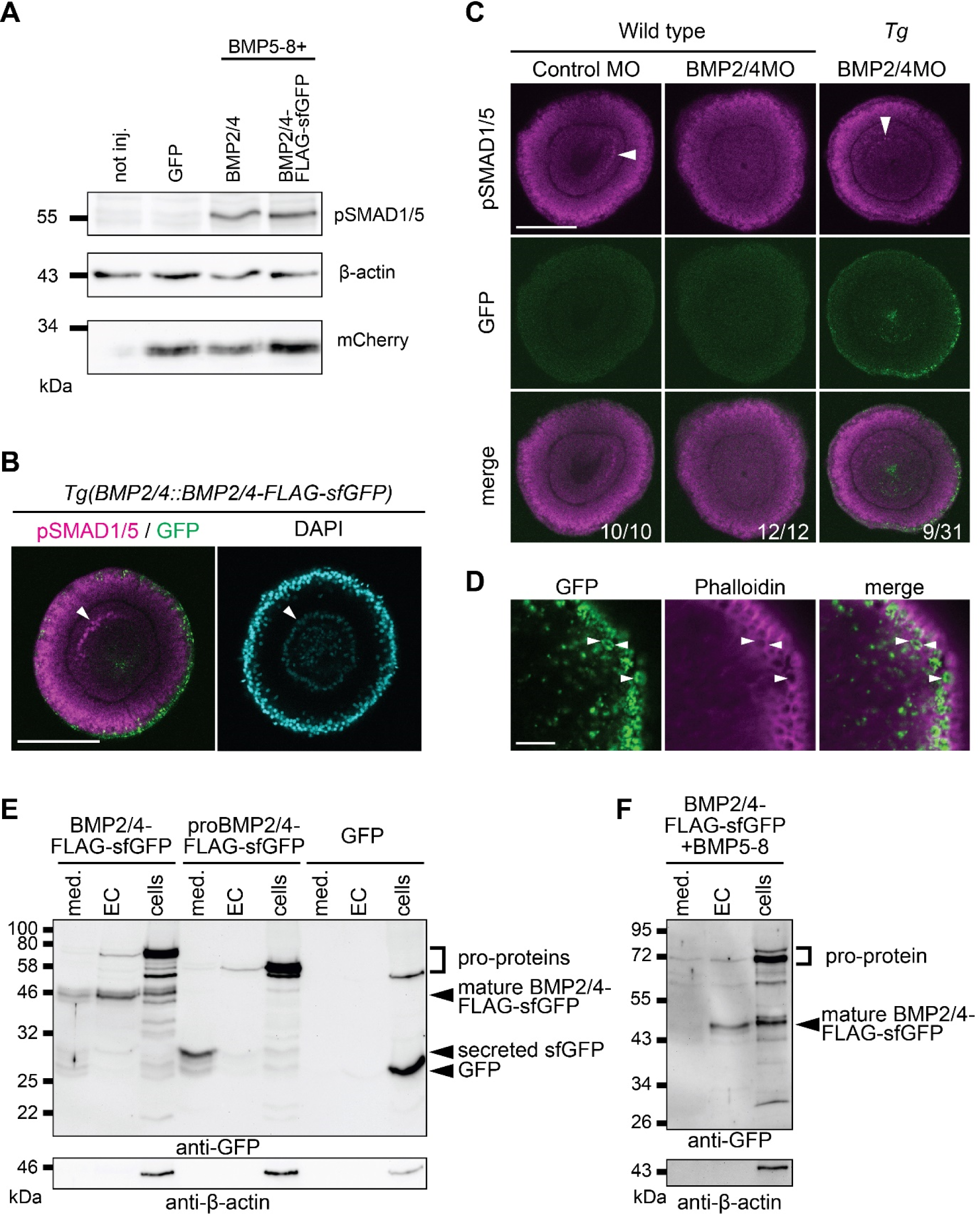
Active Nematostella BMP2/4 is retained in association with the cell surface. **A)** Anti-pSMAD1/5 western blot confirms that BMP2/4-FLAG-sfGFP can activate BMP signaling, likely as a heterodimer with BMP5-8. *GFP* mRNA was injected at equimolar concentration to the BMP mRNAs and *mCherry-CAAX* mRNA was injected at a fixed concentration in all samples as a reference. **B)** pSMAD1/5 immunofluorescence of *BMP2/4::BMP2/4-FLAG-sfGFP* embryos shows asymmetric BMP signaling, opposite to the BMP2/4 source. **C)** Anti-pSMAD1/5 immunofluorescence shows that GFP-positive *BMP2/4::BMP2/4-FLAG-sfGFP* embryos retain asymmetric BMP signaling upon injection of BMP2/4MO in 9/31 embryos. **D)** Imaging of fixed GFP-positive *BMP2/4::BMP2/4-FLAG-sfGFP* embryos shows that BMP2/4-FLAG-sfGFP signal is detected inside of the cells, close to their apical or apico-lateral surfaces. **E-F)** Western blot of medium (med.), extracellular (EC) and cellular (cells) protein fractions from embryos injected with either *BMP2/4-FLAG-sfGFP*, *proBMP2/4-FLAG-sfGFP* (control construct lacking BMP2/4 ligand domain) or *GFP* mRNA. The BMP pro-proteins are detected mostly in the cellular fractions, whereas the mature BMP2/4-FLAG-sfGFP ligand is enriched in the EC fraction. The secreted FLAG-sfGFP is detected almost exclusively in the medium. Panel (F) shows the retention of the BMP2/4-FLAG-sfGFP mature ligand in the EC fraction also when co-injected together with *BMP5-8* mRNA. Scale bars on (B-C) 100 µm. Scale bar on (D) 10 µm.

Having confirmed that BMP2/4-FLAG-sfGFP expressed from a transgene in the endogenous domain has, at least to some degree the axis-generating activity of the endogenous BMP2/4, we analyzed the localization of the BMP2/4-FLAG-sfGFP protein. The sfGFP signal was mostly detected in clusters at the surface of the embryo, indicating that BMP ligands are likely to be secreted towards the apical or apico-lateral sides of the producing cells (Fig. 2D). No extracellular BMP2/4-FLAG-sfGFP was detectable in fixed or live embryos. We concluded that the clusters of BMP2/4-FLAG-sfGFP signal were inside the producing cells, while mature, secreted BMP2/4-FLAG-sfGFP escaped detection with our methods (Supplementary Figure 1D). To obtain at least some information about the localization of the active BMP ligands, we established a fractionation protocol allowing detection of BMP ligands in the cells, on the cell surface, and in the medium by western blotting with an anti-GFP antibody. As expected, and in line with our confocal imaging data (Fig. 2D; Supplementary Fig. 1D), the bulk of the signal is observed in the cellular fraction and corresponds to unprocessed BMP2/4-FLAG-sfGFP (“cells” in Fig. 2E). Interestingly, while mature BMP ligands are secreted, they appear to remain associated with the cell surface and are mainly detected in the extracellular fraction (“EC” in Fig. 2E). In contrast, a secreted sfGFP control containing the BMP2/4 prodomain followed by sfGFP and lacking the BMP2/4 ligand domain is predominantly secreted into the medium (“med” in Fig. 2E), while cytoplasmic GFP remains inside the cells. This shows that the mature BMP2/4 ligand domain leads to retention of the BMP2/4-FLAG-sfGFP on the cell surface and suggests interactions with the extracellular matrix components or BMP receptor complexes. Similar results are obtained when BMP5-8 is co-expressed with the BMP2/4 constructs (Fig. 1F).

### Chordin does not have a strong effect on BMP stability

In the shuttling model, Chordin has pro-BMP effects that could be explained through two principle molecular mechanisms: enhanced BMP stability or enhanced BMP mobility (Mizutani et al., 2005; Müller et al., 2013). To understand whether Chordin affects BMP stability in *Nematostella*, we measured the degradation kinetics of extracellular BMP ligands. To this end, we quantified the amount of mature BMP2/4-FLAG-sfGFP ligand in the EC fraction after blocking translation with cycloheximide (Supplementary Fig. 2A-B) in two experimental conditions: “+ Chordin” (*Chd* mRNA + Control MO + *BMP2/4-FLAG-sfGFP* mRNA + *BMP5-8* mRNA + *mCherry-CAAX* mRNA) and “-Chordin” (*GFP* mRNA + ChdMO + *BMP2/4-FLAG-sfGFP* mRNA + *BMP5-8* mRNA + *mCherry-CAAX* mRNA). The anti-GFP signal from BMP2/4-FLAG-sfGFP was normalized to the mCherry signal and fitted with an exponential decay function to describe first-order degradation dynamics of the BMP2/4-FLAG-sfGFP ligand (Supplementary Fig. 2C-E). On average, extracellular BMP2/4-FLAG-sfGFP levels were higher in the absence of Chordin. The fits to the mean extracellular BMP2/4-FLAG-sfGFP levels suggest that the decay rate in the “+ Chordin” condition (*k_deg+_*=9.00·10^-5^s^-1^) is roughly twice that of the “-Chordin sample” (*k_deg-_*=4.55·10^-5^s^-1^; Figure 2B), with half-lives of 2.1 h and 4.2 h, respectively (Supplementary Fig. 2D). The observed effects are rather mild, considering the high variability between replicates, suggesting that Chordin does not have a major function in stabilizing or destabilizing BMP ligands in the “EC” fraction.

### Long-range BMP signaling requires diffusible Chordin

The main difference between the local inhibition model and the shuttling model is the necessity of Chordin mobility for Chordin-mediated BMP shuttling (Ashe and Levine, 1999; Mizutani et al., 2005; Pomreinke et al., 2017; Tuazon et al., 2020; Zinski et al., 2017). Our live imaging approaches did not reveal an extracellular BMP2/4-FLAG-sfGFP signal, hampering *in vivo* measurements of BMP mobility. Instead, we exploited the fact that no pSMAD1/5 is detectable in Chordin morphants (Genikhovich et al., 2015; Leclère and Rentzsch, 2014) to address the effect of Chordin on BMP signaling and test the requirement for extracellular Chordin mobility. Specifically, we aimed to generate a localized source of either wild-type Chordin or immobile, membrane-tethered Chordin-CD2 (Ashe and Levine, 1999; Tuazon et al., 2020) in the Chordin morphant background and use pSMAD1/5 staining as a readout of the BMP signaling activity. First, we used anti-GFP co-immunoprecipitation to confirm that i) both Chordin and Chordin-CD2 were capable of binding BMP2/4-FLAG-sfGFP, and that ii) the complex of wild-type Chordin and BMP2/4-FLAG-sfGFP was detectable in the extracellular fraction (Supplementary Fig. 3A-B). Then, we tested the biological activity of Chordin and Chordin-CD2 by injecting mRNA into zygotes and analyzing BMP signaling and *BMP2/4* expression in late gastrula stage embryos (Fig. 3A-D; Supplementary Fig. 3C-D). As expected, wild-type *Chordin* mRNA injection abolished the pSMAD1/5 gradient and radialized the expression of *BMP2/4* (Fig. 3A-B; Supplementary Fig. 3C). In contrast, bilateral symmetry was not abolished upon injection of the *Chordin-CD2* mRNA, however, pSMAD1/5 intensity was strongly reduced (Fig. 3C; Supplementary Fig. 3C). Reduced anti-BMP activity of Chordin-CD2 was not due to it having a bulky tag at the C-terminus. Chordin tagged C-terminally with FLAG-sfGFP was capable of radializing BMP2/4 expression nearly as efficiently as the wild-type Chordin; however, when we tethered such Chordin-FLAG-sfGFP to the membrane by adding the CD2 sequence to the C-terminus of sfGFP, the injected embryos developed bilateral symmetry (Supplementary Fig. 3C). We concluded that both Chordin and Chordin-CD2 were functional, and the difference in the Chordin and Chordin-CD2 overexpression phenotypes was possibly due to the presence of endogenous Chordin, which was incompletely outcompeted by Chordin-CD2. Therefore, we moved on and used Chordin-CD2 in the localized source experiments.

**Figure 3:**
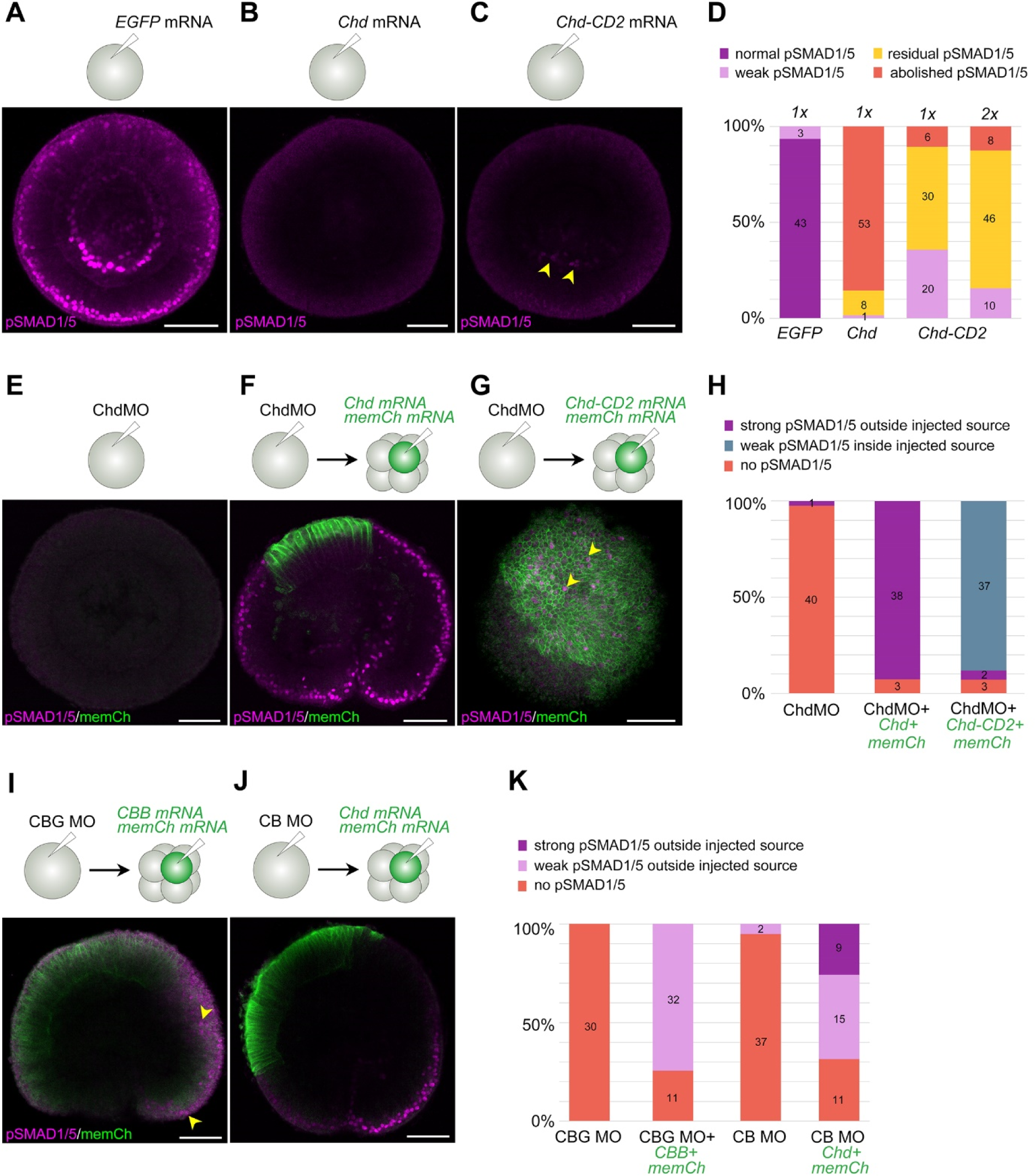
Local source experiments show that mobile Chordin shuttles BMPs. **A-D)** pSMAD1/5 immunofluorescence after injection of *GFP* (A), *Chordin* (B) and *Chordin-CD2* (C) mRNAs shows that Chordin and immobile Chordin-CD2 repress BMP signaling, although residual pSMAD1/5 signal remains in a large fraction of embryos injected with Chordin-CD2 mRNA even when the mRNA concentration is doubled (2x in D). **E)** pSMAD1/5 immunofluorescence shows absence of BMP signaling when ChdMO is injected at the one-cell stage. **F)** A local source of Chordin generated by single-blastomere mRNA injection into a Chordin morphant results in strong BMP signaling (pSMAD1/5 immunofluorescence) outside of the source cells. **G)** Same experiment with Chordin-CD2 results in weak BMP signaling inside of the source cells. **I-K)** Single blastomere injections followed by pSMAD1/5 immunofluorescence show that a source of Chordin + BMP2/4 + BMP5-8 (CBB) can activate BMP signaling at a distance in embryos that were injected with morpholinos against *Chordin*, *BMP2/4* and *GDF5-like* (CBG MO in I). GDF5-like is therefore not required to activate BMP signaling in non-source cells. In embryos injected with ChdMO + BMP2/4MO (CB MO), a local source of Chordin is sufficient to trigger BMP signaling outside of the source, indicating that Chordin promotes also GDF5-like-mediated signaling (J). Scale bars 50 µm.

Previously we showed that the expression of all three *Nematostella* BMP ligand genes active in the early embryo, *BMP2/4*, *BMP5-8* and *GDF5-like*, is nearly or completely abolished at 2 dpf upon Chordin knockdown (Genikhovich et al., 2015). However, we knew that at 1 dpf (late gastrula) they were still active, although their expression was radialized (Supplementary Fig. 3D). Thus, we concluded that BMP ligands are initially present in Chordin morphants and analyzed BMP signaling activity in Chordin morphants injected into a single cell at the 8-cell stage with mRNAs of either wild-type Chordin or membrane-tethered, immobile Chordin-CD2 (*Chd* and *Chd-CD2* in Fig. 3, respectively). To label the source of Chordin, *mCherry-CAAX* (*memCh* in Fig. 3) mRNA was co-injected with the *Chordin* mRNAs (Fig. 3E-H). Chordin morpholino injection (ChdMO in Fig. 3) abolished BMP signaling in late gastrula (1 dpf) despite the presence of the BMPs (Fig. 3E), suggesting that BMP signaling in *Nematostella* is Chordin-dependent at this stage. In contrast, creating a local memCherry-labelled source of either Chordin or Chordin-CD2 rescued BMP signaling in Chordin morphants, however, in a strikingly different manner. In Chordin morphants with a local source of wild-type Chordin, we observed strongly stained pSMAD1/5-positive nuclei outside the Chordin source (Fig. 3F), while in Chordin morphants with a local source of Chordin-CD2, we saw weakly stained pSMAD1/5-positive nuclei inside the Chordin-CD2 source (Fig. 3G). Moreover, upon simultaneous morpholino knockdown of BMP2/4, GDF5-like and Chordin (CBG MO in Fig. 3I,K), which abolishes not only Chordin but also the endogenous BMPs, a local source of Chordin, BMP2/4 and BMP5-8 (CBB mRNA in Fig. 3I and 3K) was sufficient to rescue BMP signaling in approximately 74% of the embryos (Fig. 3I and 3K, Supplementary Fig. 3E). We reasoned that injection of BMP5-8MO was unnecessary in this case, since the BMP2/4MO and BMP5-8MO phenotypes are identical, most likely due to the abovementioned putative obligatory heterodimerization of these two BMPs. Finally, we tested whether Chordin was capable of promoting only BMP2/4/BMP5-8-mediated signaling or whether it could do the same with GDF5-like-mediated signaling. *GDF5-like* mRNA is present in 1 dpf embryos both upon *Chd* and *BMP2/4* knockdown (Supplementary Fig. 3D), therefore we injected zygotes with a mixture of Chordin and BMP2/4 morpholinos without the GDF5-like morpholino (CB MO in Fig. 3J and 3K) and created local sources of Chordin but did not introduce any exogenous BMPs. In such embryos, BMP signaling on the opposite side to the Chordin source was rescued in approximately 69% of the cases, moreover, in about 26% of the cases BMP signaling was strong (Fig. 3J and 3K, Supplementary Fig. 3E). In summary, we conclude that extracellular Chordin mobility is required for long-range activation of BMP signaling, and that the agonistic action of Chordin and BMP is not selective with regard to the type of BMP ligand.

## Discussion

Chordin-dependent BMP signaling is required to establish and pattern the second, directive body axis in the bilaterally symmetric non-bilaterian, the sea anemone *Nematostella vectensis* (Genikhovich et al., 2015; Leclère and Rentzsch, 2014). However, although “BMP shuttling” has been suggested to be the mechanism underlying this process (Genikhovich et al., 2015), a “local inhibition” model may be a valid alternative. Here, we attempted to learn more about the molecular mechanism of *Nematostella* directive axis formation and the role of Chordin in it.

### BMP heterodimer transport appears to occur in an unexpected location

Our current understanding of BMP signaling in *Nematostella* directive axis patterning is based on a model, in which BMP heterodimers containing BMP2/4 and BMP5-8 diffuse through the mesoglea – a layer of extracellular matrix separating the outer ectodermal layer from the inner endoderm and mesoderm (Genikhovich et al., 2015). We showed that injection of a mixture of *BMP2/4* and *BMP5-8* mRNAs indeed elicits a stronger pSMAD1/5 signal than overexpression of these ligands individually. This is consistent with earlier loss-of-function data (Genikhovich et al., 2015; Leclère and Rentzsch, 2014; Saina et al., 2009) and suggests that, BMP2/4/BMP5-8 heterodimers are the biologically relevant BMP ligands during axis patterning in *Nematostella*. This is similar to the situation in Bilateria, where BMP heterodimers outperform homodimers in various developmental contexts (Bauer et al., 2023; Little and Mullins, 2009; Tajer et al., 2021). However, we still do not understand where BMP diffusion and signaling takes place. Although our attempts to visualize extracellular sfGFP-tagged BMP2/4 failed, we made two important observations. First, the visualization of the intracellular sfGFP-tagged BMP2/4 suggested that it was secreted towards the apical or apico-lateral surface of the ectodermal cells of the *Nematostella* gastrula rather than basally, towards the mesoglea. Second, in our fractionation experiments, we showed that mature BMP ligands were retained on the surface of the cells rather than released into the medium.

The potentially apical or apico-lateral secretion of BMPs is consistent with a previously observed apico-lateral localization of the ectopically expressed GFP-tagged constitutively active type I BMP receptor Alk6 (Genikhovich et al., 2015). Secretion towards the outside of the embryo, if experimentally confirmed in future studies, would also not be a unique feature of *Nematostella*. In *Drosophila*, BMP2/4 also appears to be secreted towards the surface of the embryo and shuttling is thought to happen in the perivitelline space, a sealed-off extraembryonic compartment (Wang and Ferguson, 2005). However, *Nematostella* embryos are not surrounded by any extraembryonic membrane and the retention of BMP ligands in the “EC” fraction observed in our fractionation experiments may be necessary to prevent the loss of the signaling molecules into the medium and facilitate gradient formation (Müller et al., 2013). Clearly, the assumptions of our 2015 model about the geometry of the BMP diffusion and the BMP signaling domain will need to be revisited once we know more about the distribution of mature BMP ligands and receptors in the embryo and the mechanism of BMP retention on the cell surface.

### BMP shuttling as a candidate ancestral mechanism of second axis patterning

BMP-dependent axial patterning systems, although extremely diverse in different animal groups (for review, see Mörsdorf et al., 2024), tend to repeatedly evolve a “seesaw” regulatory architecture with both ends of the second body axis expressing different BMP ligands and BMP signaling occurring on one of these two ends, opposite to the Chordin expression domain. There are exceptions, especially in insects, where BMPs signal dorsally and Toll signaling plays a role as a ventral signal (Arora et al., 1994; Ferguson and Anderson, 1992; Nunes da Fonseca et al., 2008; Özüak et al., 2014; Sachs et al., 2015; van der Zee et al., 2006), however, in anthozoan Cnidaria, such as *Nematostella*, and in Deuterostomia this seems to be the general rule. For example, in the frog *Xenopus*, the “ventral signaling center” expresses BMP4 and BMP7, the “dorsal signaling center” expresses BMP2, ADMP (another BMP ligand) and Chordin, and pSMAD1/5 gradient has a ventral maximum (Plouhinec et al., 2013). In the sea urchin *Paracentrotus*, the “ventral signaling center” expresses BMP2/4, ADMP1 and Chordin, the “dorsal signaling center” expresses ADMP2, and the pSMAD1/5 gradient has a dorsal maximum (Lapraz et al., 2009; Lapraz et al., 2015). In both situations, Chordin-mediated BMP shuttling has been suggested as a patterning mechanism (Ben-Zvi et al., 2008; Lapraz et al., 2015; Plouhinec et al., 2013). The expression of BMP2/4, BMP5-8 and Chordin on one side of the directive axis, of GDF5-like on the other side of the directive axis, and a pSMAD1/5 gradient maximum on the GDF5-like-expressing side in the embryos of the cnidarian *Nematostella* fits neatly to the seesaw paradigm. One unique feature of the *Nematostella* BMP-dependent axial patterning is the extent to which it relies on Chordin in 1-2 dpf embryos. In all bilaterian models, where this has been addressed, Chordin loss-of-function de-represses BMP signaling expanding the pSMAD1/5-positive domain (Lapraz et al., 2009; Peluso et al., 2011; Plouhinec et al., 2013; Pomreinke et al., 2017; Tan et al., 2022; Zinski et al., 2017). In contrast, Chordin knockdown leads to the disappearance of the pSMAD1/5-positive domain in *Nematostella*, despite the presence of *BMP2/4*, *BMP5-8* and *GDF5-like* mRNA in the embryo at gastrulation (Genikhovich et al., 2015; Leclère and Rentzsch, 2014; Fig. 3B; Supplementary Fig. 3D). In light of this striking pro-BMP effect of *Nematostella* Chordin it was important to verify that it can still act as a BMP antagonist, which we could confirm in our overexpression experiments both with the wild-type Chordin and with the membrane-tethered Chordin-CD2.

Shuttling of BMP2/4 and BMP5-8 ligands by Chordin from the “low BMP signaling” to the “high BMP signaling” of the directive axis provided a plausible explanation for the loss-of-function phenotypes observed in *Nematostella*, however, an alternative “local inhibition” mechanism described in zebrafish, in which Chordin is simply a local BMP repressor, could not be excluded (Genikhovich et al., 2015; Pomreinke et al., 2017; Tuazon et al., 2020; Zinski et al., 2017). To find out whether Chordin acts as a local BMP inhibitor or a long-range BMP shuttle during the directive axis formation in *Nematostella*, we established a localized source assay. We showed that mobile Chordin was required to promote strong BMP signaling at the side of the embryo opposite to the Chordin source, in line with the shuttling model. In contrast, not only was our membrane-tethered Chordin-CD2 unable to activate BMP signaling at a distance, but it activated it inside the Chordin-CD2 source instead. While this apparent local pro-BMP effect is surprising for a BMP inhibitor like Chordin, such behavior is not unheard of. Crossveinless-2 (CV2) is a BMP-binding protein with Chordin type cysteine-rich (CR) domains, which attaches to the cell surface by interacting with the side chains of heparan sulphate proteoglycans. Both, in *Drosophila* and in vertebrates, CV2 is positively regulated by BMP signaling and exhibits complex pro- and anti-BMP effects (Ambrosio et al., 2008; Ikeya et al., 2006; Rentzsch et al., 2006; Serpe et al., 2008). In *Nematostella*, a CV2 ortholog is not expressed in 1-2 dpf embryos, but becomes detectable in the pSMAD1/5 positive domain later in development (Fig. 4A-C). Importantly, in *Drosophila*, the pro-BMP function of CV-2 was suggested to be due to it “handing over” sequestered BMPs to the type I BMP receptor Thickveins (Serpe et al., 2008). We speculate that by adding the CD2 sequence to the C-terminus of Chordin we may have created a CV2 analogue. Its behavior indicates the direction for future research: it will be important to test experimentally whether wild-type, mobile Chordin facilitates the BMP-receptor interaction in 1-2 dpf *Nematostella* embryos. Should this be the case, this would explain the loss of BMP signaling in Chordin morphants despite the presence of BMPs in the embryo. On the other hand, in 4 dpf embryos, when the directive axis is already patterned, Chordin expression stops, and BMP signaling becomes confined to the areas where BMPs are expressed (Knabl et al., 2024). Given the indispensability of Chordin for BMP signaling in the 1-2 dpf embryos, it will be important to find out how it is possible that Chordin becomes unnecessary during later stages.

**Figure 4:**
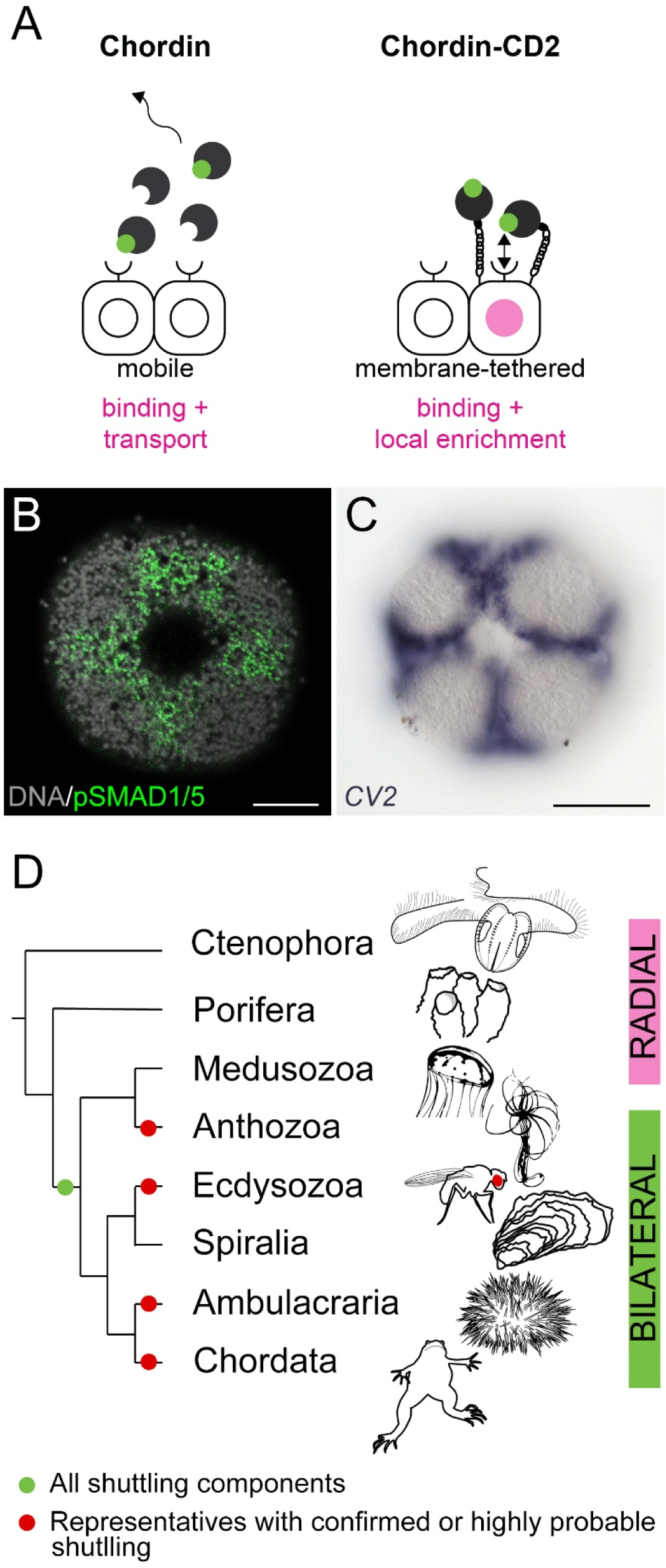
BMP shuttling-mediated bilaterality may have been present in the cnidarian-bilaterian ancestor. **A)** BMP binding by the diffusible wild-type Chordin results in BMP ligand transport and local inhibition of BMP signaling. BMP binding by the membrane-tethered Chordin-CD2 results in a weak local activation of BMP signaling, potentially due to an enrichment of the BMP ligands close to the cell surface and a stimulation of the BMP-receptor interaction. **B)** BMP signaling activity in the oral ectoderm of the 4 dpf late planula of Nematostella. pSMAD1/5 activity is observed between the future primary tentacle buds and around the mouth. **C)** Expression of *Nematostella CV2* in the same domain. **D)** Phylogenetic distribution of Chordin-mediated BMP shuttling components across Metazoa. Scale bars on (B-C) 50 µm.

## Conclusion

Our work shows that Chordin-dependent BMP shuttling described in Bilateria is indeed the mechanism responsible for the establishment and patterning the second body axis in the sea anemone *Nematostella*, a member of the bilaterian sister clade. Thus, the last common ancestor of Cnidaria and Bilateria had all the necessary components to establish a BMP shuttling-mediated symmetry break and maintain the second body axis (Fig 4D). Whether this means that the cnidarian-bilaterian ancestor was a bilaterally symmetric animal we cannot be certain, but it is quite possible. On the other hand, the combination of BMP ligands and Chordin represents a powerful system to break symmetries and drive axial patterning, which in different animals has been assembled into such a variety of regulatory networks with different topologies, that convergent evolution of bilaterality in Cnidaria and Bilateria cannot be excluded.

## Materials and methods

### Fusion constructs, in vitro transcription and transgenics

The sequences for expression constructs were generated using standard cloning techniques, the primers that were used are shown in the Supplementary Table 1. Fusion constructs were generated via splicing by overlap extension PCR (Heckman and Pease, 2007). In BMP2/4-FLAG-sfGFP, the FLAG-sfGFP tag is placed between the BMP2/4 pro-domain and the mature ligand domain after the predicted cleavage site RRKRSL (Duckert et al., 2004) with LGDPPVAT linkers, mimicking previous zebrafish constructs (Pomreinke et al., 2017). The BMP2/4 coding sequence in the construct contains four silent mutations to block binding of the previously tested BMP2/4MO (Saina et al., 2009). BMP2/4-mCherry is an equivalent fusion construct where mCherry replaces FLAG-sfGFP. proBMP2/4-FLAG-sfGFP corresponds to BMP2/4-FLAG-sfGFP with the mature ligand domain removed. Chordin-FLAG-sfGFP encodes a protein where FLAG-sfGFP is fused to the C-terminus of Chordin. For Chordin-CD2 and Chordin-FLAG-sfGFP-CD2, the CD2 sequence (Ashe and Levine, 1999) is fused to the C-terminus of Chordin and Chordin-FLAG-sfGFP, respectively. mRNA coding for mCherry with the C-terminal CAAX sequence (Choy et al., 1999) was used to label the mRNA injected cells in the localized source experiments. Sequences for mRNA synthesis were cloned downstream of the T7 promoter into a customized pJet1.2 plasmid (Thermo Fisher) carrying PacI and SbfI sites followed by the SV40 polyadenylation signal. The mMessage mMachine T7 Transcription Kit (Thermo Fisher) was used for in vitro transcription. To generate the transgenic BMP2/4::BMP2/4-FLAG-sfGFP line, a 4.5 kb DNA fragment upstream of the translation start site of the *BMP2/4* gene (Schwaiger et al., 2014) was cloned into a previously described transgenesis vector (Renfer and Technau, 2017) to drive the expression of *bmp2/4-FLAG-sfGFP*. 50 ng/µl plasmid was digested with 0.2 U I-SceI/µl for 30 min at 37°C before injection. For live-imaging of the *BMP2/4::BMP2/4-FLAG-sfGFP* embryos, an F0 transgenic male was crossed to a WT female, and the embryos were injected at the one-cell stage with 61 ng/µl *mCherry-CAAX* mRNA. GFP-positive embryos were mounted in 2% low-melting agarose in *Nematostella* medium on a cover slip. Upon solidification of the agarose, the cover slip was placed in a Petri dish lid, covered with *Nematostella* medium and imaged on a Leica TCS SP8 confocal microscope using a 40x water-dipping objective.

### Gene knockdown, overexpression and the local source assay

Previously characterized morpholino oligonucleotides against *Chd*, *BMP2/4* and *GDF5-like*, (Genikhovich et al., 2015; Saina et al., 2009) were injected into dejellied *Nematostella* zygotes as described previously (Renfer and Technau, 2017). A splice MO targeting exon 9 of *Mef2* was used as a Control MO as in (Kraus et al., 2016), since it does not affect development at least until primary polyp stage and shows lower toxicity than the Gene Tools standard control MO. All MO sequences can be found in the Supplementary Table 2. For overexpression, mRNAs were injected into zygotes at concentrations summarized in Supplementary Table 3. All injection mixes contained 30 ng/µl Dextran-AlexaFluor568 (Invitrogen) as a tracer. For the local source assay, *Nematostella* zygotes were injected with the splice morpholino against Chordin (ChdMO) with or without BMP2/4MO and GDF5lMO and kept at 18.5°C until they started dividing. When the embryos reached 8-cell stage, individual blastomeres were injected with mRNA mixtures described in the Supplementary Table 3. The embryos were kept at 21°C and fixed at 1 dpf (24-26 hours post-fertilization, hpf) for anti-pSMAD1/5 and anti-mCherry immunostaining as described below.

### Immunohistochemistry, in situ hybridization

For immunohistochemistry and in situ hybridization, the embryos were fixed in ice-cold 0.25% glutaraldehyde/3.7% formaldehyde/PTx (PTx=1x PBS with 0.3% Triton X100) for 2 minutes on ice and then in 3.7% formaldehyde/PTx for 1 hour at 4°C with overhead rotation. For immunohistochemistry, fixed embryos were washed five times for 5 minutes in PTx, then incubated in pre-chilled methanol on ice for 8 minutes, washed three more times with PTx, and blocked in a blocking solution containing 5% heat-inactivated sheep serum and 95% of 1% BSA/PTx for 2 hours at room temperature. At the same time, primary antibodies were diluted and pre-incubated in the blocking solution. The embryos were stained overnight with rabbit monoclonal anti-pSMAD1/5/9 (Cell Signaling, 13820) in 1:200 dilution and mouse monoclonal anti-mCherry antibody (Takara, 632543) in a 1:400 dilution at 4°C on rocker. After five15-minute PTx washes the embryos were blocked again, and primary antibodies were detected with goat anti-rabbit AlexaFluor 488 (Invitrogen, A-11008) and goat anti-mouse AlexaFluor 633 (Invitrogen, A-21050) antibodies in a 1:1000 dilution for 2 hours at room temperature on the rocker. 5 µg/ml DAPI was added to the secondary antibody solution to make sure that pSMAD1/5 signal is nuclear. sfGFP fluorescence was visible after fixation and no additional antibody staining was performed; AlexaFluor633-phalloidin (Invitrogen, A22284) was used at a final concentration of 4 units/ml. The embryos were imaged with a Leica SP8 or Leica Stellaris 5 LSCM.

For in situ hybridization, the embryos were fixed as described above, washed five times in PTx or PTw (PTw=1xPBS with 0.1% Tween 20) and once in 100% methanol, and stored in 100% methanol at −20°C. After rehydration by washing for 5 minutes in 50% methanol/PTw and in pure PTw, the embryos were handled as described in Kraus et al., 2016 with the following change: proteinase K treatment was performed with 10 µg/ml proteinase K/PTw solution for 20 minutes.

### Embryo fractionation to sample secreted proteins

Embryos were injected with the different combinations of mRNAs (equimolar between samples) and incubated at 21°C until the cells started dividing. 100 dividing embryos of a sample were transferred to a well of a 96-well plate in 90 µl *Nematostella* medium and incubated at 21°C overnight. Around 22 hpf, the embryos and the medium were transferred to a 1.5 ml reaction tube. 80 µl medium were removed and mixed with 25 µl 5x loading dye as “medium” fraction. 80 µl Mg^2+^/Ca^2+^-free artificial sea water (27 g/L NaCl, 1 g/L Na_2_SO_4_, 0.8 g/L KCI, 0.18 g/L NaHCO_3_; Kirillova et al., 2018) containing cOmplete Protease Inhibitor Cocktail (Roche) were added to the embryos, and embryos were dissociated into a cell suspension by trituration. After 1 min centrifugation at 1000 rcf (RT), 80 µl supernatant were mixed with 25 µl 5x gel loading dye as “EC” (extracellular) fraction. The cells were then lysed in 80 µl cold Cell Extraction Buffer (Life Technologies/Invitrogen) containing cOmplete Protease Inhibitor Cocktail (Roche). After 10 min centrifugation at 16000 rcf (4°C), 80 µl supernatant were collected and mixed with 25 µl 5x gel loading dye as “cells” fraction.

### Co-immunoprecipitation

Embryos were injected with a mix of *BMP5-8* and *BMP2/4-mCherry* mRNAs containing either *Chordin* mRNA, *Chordin-CD2* mRNA or *GFP* mRNA (negative control). At 21 hpf, 230 embryos per sample were dissociated in 130 µl Mg^2+^/Ca^2+^-free artificial sea water (with protease inhibitors) to prepare EC protein fractions as described above. The cells were lysed in 130 µl lysis buffer (10 mM TRIS, 150mM NaCl, 0.5 mM EDTA, 0.5% NP40 substitute (Merck); pH 7.5) containing cOmplete Protease Inhibitor Cocktail (Roche). For the immunoprecipitation (IP), GFP-Trap Magnetic Agarose beads (Chromotek) were used according to the manufacturer’s recommendations. Per IP, 12.5 µl agarose slurry were used and equilibrated by rinsing with 500 µl dilution buffer (10 mM TRIS, 150mM NaCl, 0.5 mM EDTA; pH 7.5) three times. The agarose was suspended in dilution buffer for IPs from cell lysates and in a 2 : 1 lysis buffer : dilution buffer mix for IPs from EC fractions and supplemented with cOmplete Protease Inhibitor Cocktail (Roche). Of the 130 µl EC fraction/cell lysate, 6 µl were saved as 5 % input and 120 µl were mixed with 180 µl beads prepared for the IP as described above. During IP, the tubes were incubated at 4°C for 1 h with overhead rotation. The beads were then washed three times for 5 min with 500 µl wash buffer (10 mM TRIS, 150 mM NaCl, 0.5 mM EDTA, 0.05% NP40 substitute (Merck); pH 7.5) containing cOmplete Protease Inhibitor Cocktail (Roche), and transferred to a fresh tube during the last wash step. To collect the IP samples, the beads were heated to 95°C in 50 µl 1x protein loading buffer for 5 min.

### Cycloheximide treatment

To optimize the efficiency of translation inhibition using cycloheximide (CHX), we performed a test experiment in which we injected embryos with 20 ng/µl HSP70::GFP plasmid, in which the 1.7 kb promoter region of *Nematostella Hsp70* was driving the expression of GFP. At 1 dpf, they were incubated in *Nematostella* medium containing 200, 500 or 1000 ng/µl CHX or 1% DMSO (corresponding to the highest CHX concentration) for 30 min. Afterwards, they were heat shocked in a water bath at 37°C for 30 min. Control embryos were kept at 21°C. Cell lysates for western blot were made 5.5 h post heat shock. This experiment (Supplementary Figure 2A-B) revealed efficient inhibition of translation with 500 ng/µl CHX and this concentration was used for all other experiments. Since translation inhibition was effective within ca. 30 min after the start of CHX incubation, 30 min after the start of CHX incubation was considered the “0 h” timepoint in the time course experiment. Cellular and extracellular protein fractions in the time course experiment were prepared from 50 embryos as described above using 20 µl Mg^2+^/Ca^2+^-free artificial sea water and 20 µl cell extraction buffer, respectively.

### SDS-PAGE and western blot

Whole-embryo protein lysates were prepared with Cell Extraction Buffer (Life Technologies/Invitrogen) containing cOmplete Protease Inhibitor Cocktail (Roche) that was additionally supplemented with PhosSTOP Phosphatase Inhibitor Cocktail (Roche) if the aim was to detect pSMAD1/5. Proteins were separated on 10 % polyacrylamide gels and blotted onto a nitrocellulose membrane at 100 V for 1 h using the Mini Trans-Blot system (Bio-Rad). Membranes were blocked with 5 % milk powder in PTw (1x PBS, 0.1 %Tween-20) and the same blocking solution was used for the following antibody dilutions: 1:10000 anti-GFP (Abcam ab290), 1:2000 monoclonal anti-mCherry (Takara/Clontech #632543), 1:1000 polyclonal anti-mCherry (Chromotek pabr1; used only for the detection of BMP2/4-mCherry in CoIP experiments), 1:10000 anti-β-Actin (Cell Signaling #4970) and 1:10000 of the HRP-conjugated Anti-Rabbit IgG (Promega W401B) and Anti-Mouse IgG (Promega W402B). For the pSMAD1/5 western blot, the protocol of (Watanabe et al., 2014) was followed using 1:1000 anti-phospho-SMAD1/SMAD5/SMAD9 (Cell Signaling #11971) and 1:10000 HRP-conjugated Anti-Rabbit IgG (Promega W401B). The SuperSignal West Femto Maximum Sensitivity Substrate (Thermo Scientific) was used for enhanced chemiluminescence detection. Western blot band intensities were quantified in Fiji (Schindelin et al., 2012) using rectangular ROIs. Background intensities were subtracted and the background-subtracted intensities normalized as described. A single-exponential decay function *(1)* was fit to the data in Supplementary Figure 2D-E using the nls() function in R (R Core Team, 2020).

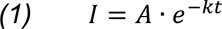

Here, *I* is the intensity, *A* is the intensity at timepoint 0, *k* is the degradation rate, *t* is time.

## Acknowledgements

We thank Hillary Ashe for providing *Drosophila sog-CD2*, Patrick Müller for sharing the *sfGFP* construct, Simon Weinberger for cloning the *BMP2/4* promoter element, Eduard Renfer for generating the *Hsp70::GFP* plasmid, Annamaria Sgromo, Paul Knabl and Sophie Frampton for discussions and feedback on the manuscript. Confocal microscopy was performed at the Core Facility Cell Imaging and Ultrastructure Research, University of Vienna - member of the Vienna Life-Science Instruments (VLSI). This research was funded in whole or in part by the Austrian Science Fund (FWF) grant (DOI 10.55776/P32705) to G.G. and by the Austrian Science Fund (FWF) Lise-Meitner Fellowship (DOI 10.55776/M3291) to D.M.. For the purpose of Open Access, the author has applied a CC BY public copyright license to any Author Accepted Manuscript (AAM) version arising from this submission.

---

**Supplementary Figure 1:**
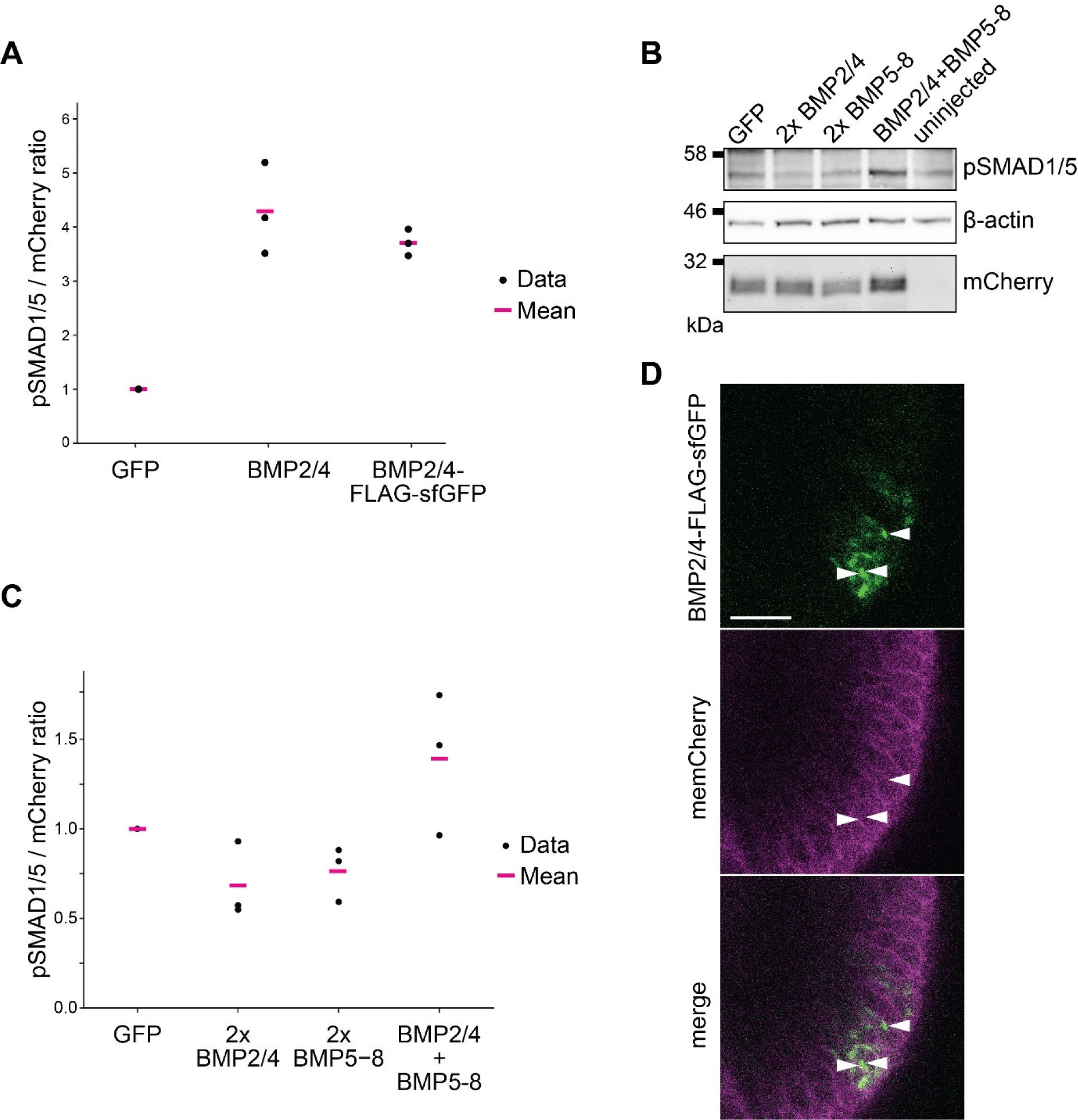
BMP2/4-FLAG-sfGFP is an active ligand eliciting stronger pSMAD1/5 signal in combination with BMP5-8. **A)** Quantification of the western blot shown in Fig. 2A and two additional independent replicates. After quantification, the pSMAD1/5-to-mCherry ratio was normalized to the GFP sample. Black dots show measurements from individual replicates, pink bars show mean values. **B)** Western blot shows that co-injection of *BMP2/4* and *BMP5-8* mRNAs elicits a stronger pSMAD1/5 signal than each mRNA alone injected at double the concentration (labelled as 2x). mCherry signal was used as an injection reference. **C)** Quantification of data in (B) and two additional, independent replicates. Black dots show values from individual measurements, pink bars show mean values. **D)** Live imaging of 1 dpf *BMP2/4::BMP2/4-FLAG-sfGFP* embryos injected with *mCherry-CAAX* mRNA (memCherry) at the one-cell stage show GFP signal inside the cells. Scale bar 20 µm.

**Supplementary Figure 2:**
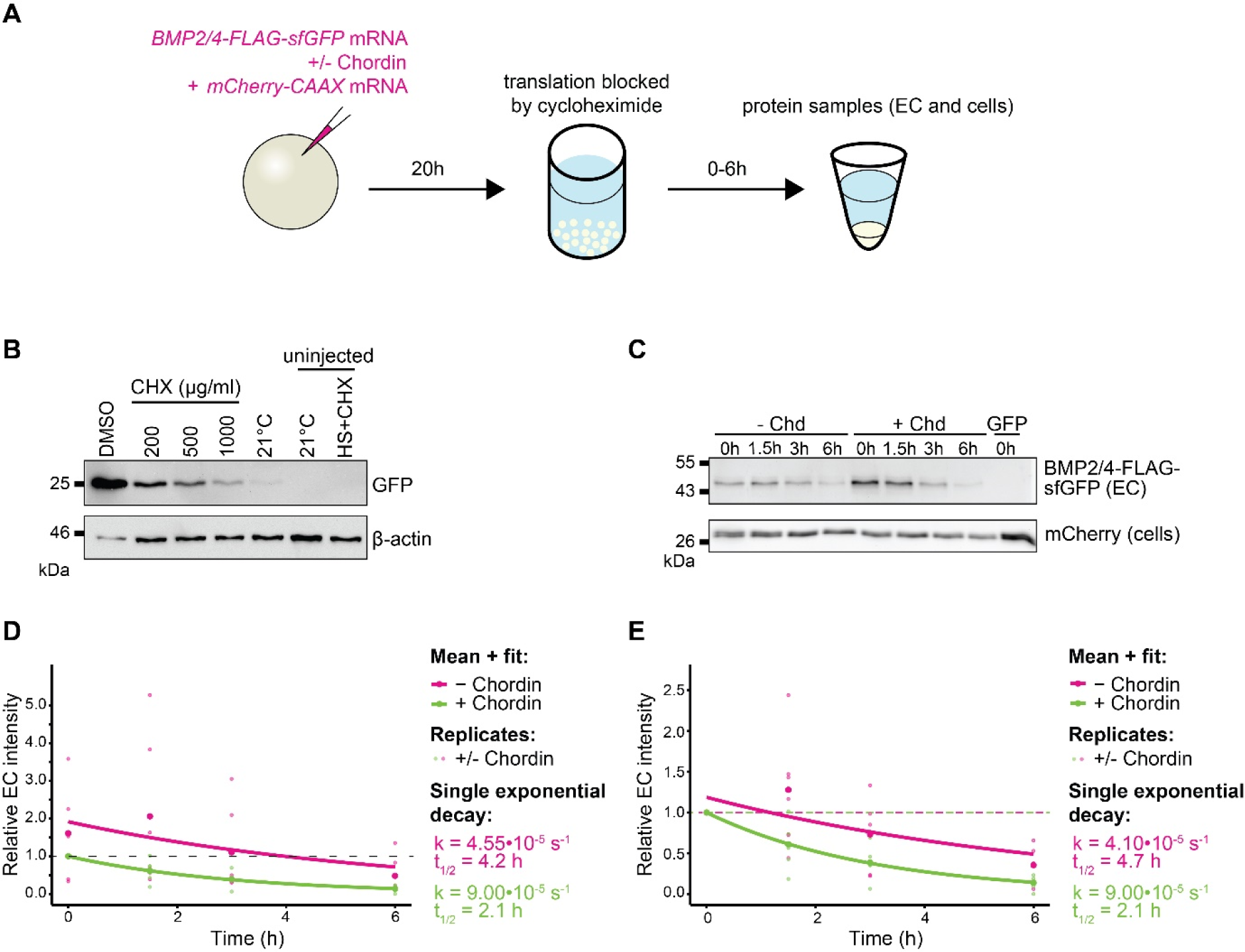
Chordin does not have a strong effect on BMP stability. **A)** A scheme of cycloheximide (CHX) treatment after mRNA and morpholino microinjection. **B)** GFP translation is efficiently blocked by CHX in the *Nematostella Hsp70::GFP* injected embryos. Anti-GFP western blot shows efficient inhibition of GFP translation in the presence of CHX. Controls were kept at 21°C. **C)** Extracellular mature BMP2/4-FLAG-sfGFP levels in the presence or absence of Chordin over 6 h time course. mCherry signal serves as injection reference. **D)** Relative EC intensity (BMP2/4-FLAG-sfGFP / mCherry ratio) of six independent measurements. Line shows single exponential decay function fit to the mean data. All data points are normalized to the 0 h point of the “+ Chordin” sample. **E)** Same experimental data as in D), but “-Chordin” and “+ Chordin” samples were normalized independently to their 0 h sample.

**Supplementary Figure 3:**
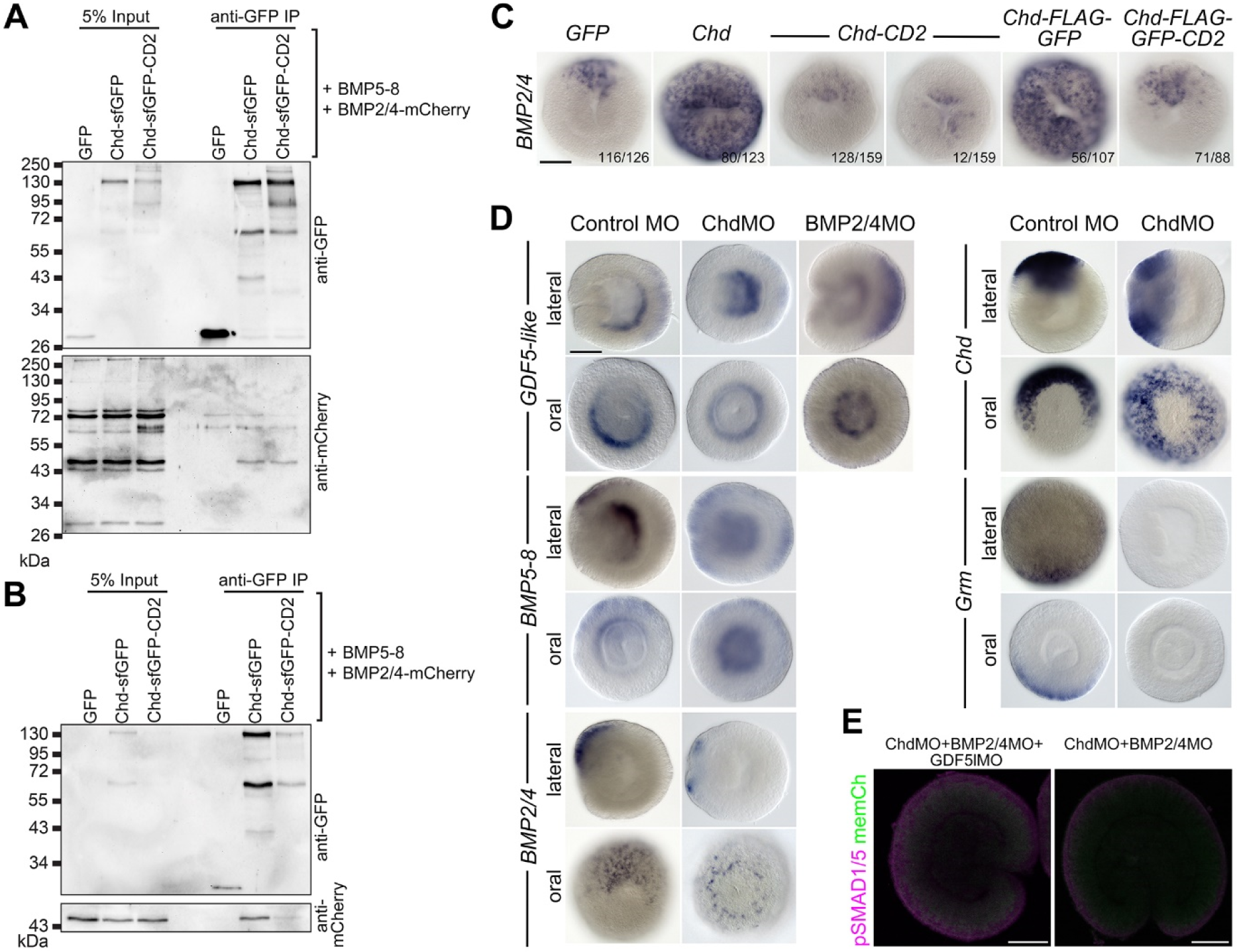
Chordin forms a complex with BMPs in the extracellular fraction and can act as a BMP inhibitor. **A)** Anti-GFP CoIP from cellular protein fractions of embryos expressing both BMP5-8 and BMP2/4-mCherry as well as one of the given bait proteins. BMP2/4-mCherry co-immunoprecipitates with both Chordin-FLAG-sfGFP (Chd-sfGFP) and Chordin-FLAG-sfGFP-CD2 (Chd-sfGFP-CD2). **B)** Anti-GFP CoIP from extracellular protein fractions of the same embryos used in A). BMP2/4-mCherry co-immunoprecipitates with Chordin-FLAG-sfGFP (Chd-sfGFP), revealing the presence of an extracellular BMP-Chordin complex. **C)** mRNA injection followed by *BMP2/4 in situ* hybridization at 30 hpf shows that Chordin and Chordin-FLAG-sfGFP repress BMP signaling, resulting in radialized BMP2/4 expression. *Chordin-CD2* and *Chordin-FLAG-sfGFP-CD2* mRNA injections result in a mostly asymmetric expression of *BMP2/4*, indicating the presence of a BMP signaling gradient. **D)** *In situ* hybridization after the injection of ChdMO shows that the BMP ligands *BMP2/4*, *BMP5-8* and *GDF5-like*, as well as *Chd* are expressed in a radially symmetric manner at 1 dpf, whereas the expression of *Grm* is lost. *GDF5-like* mRNA is present in a radially symmetric domain in embryos injected with BMP2/4MO. **E)** Similar to ChdMO injection, combined knockdown of Chd, BMP2/4 and GDF5-like or Chd and BMP2/4 completely abolishes BMP signaling activity. Scale bars on (C-E) 50 µm.

**Supplementary Table 1:**
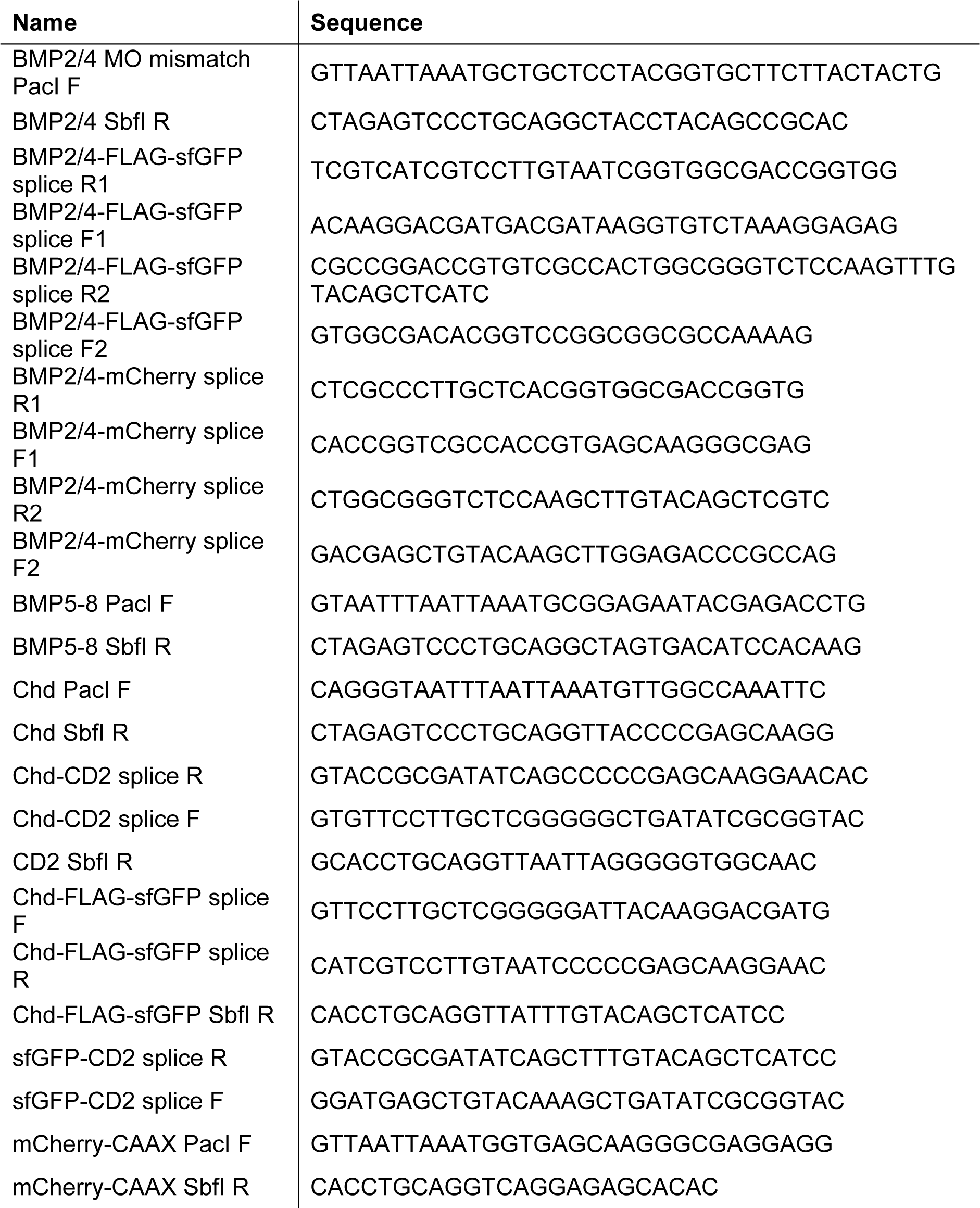
Primers used in this study.

**Supplementary Table 2:**
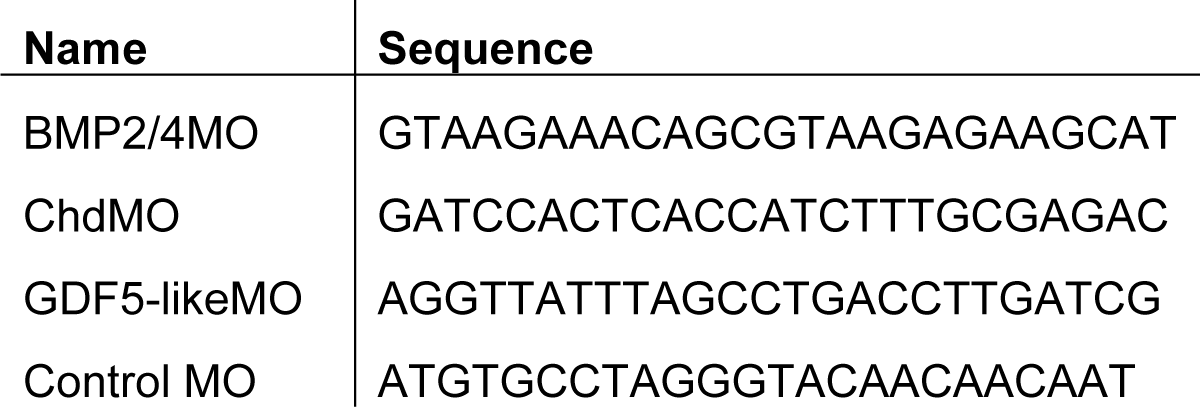
MOs used in this study.

**Supplementary Table 3:**
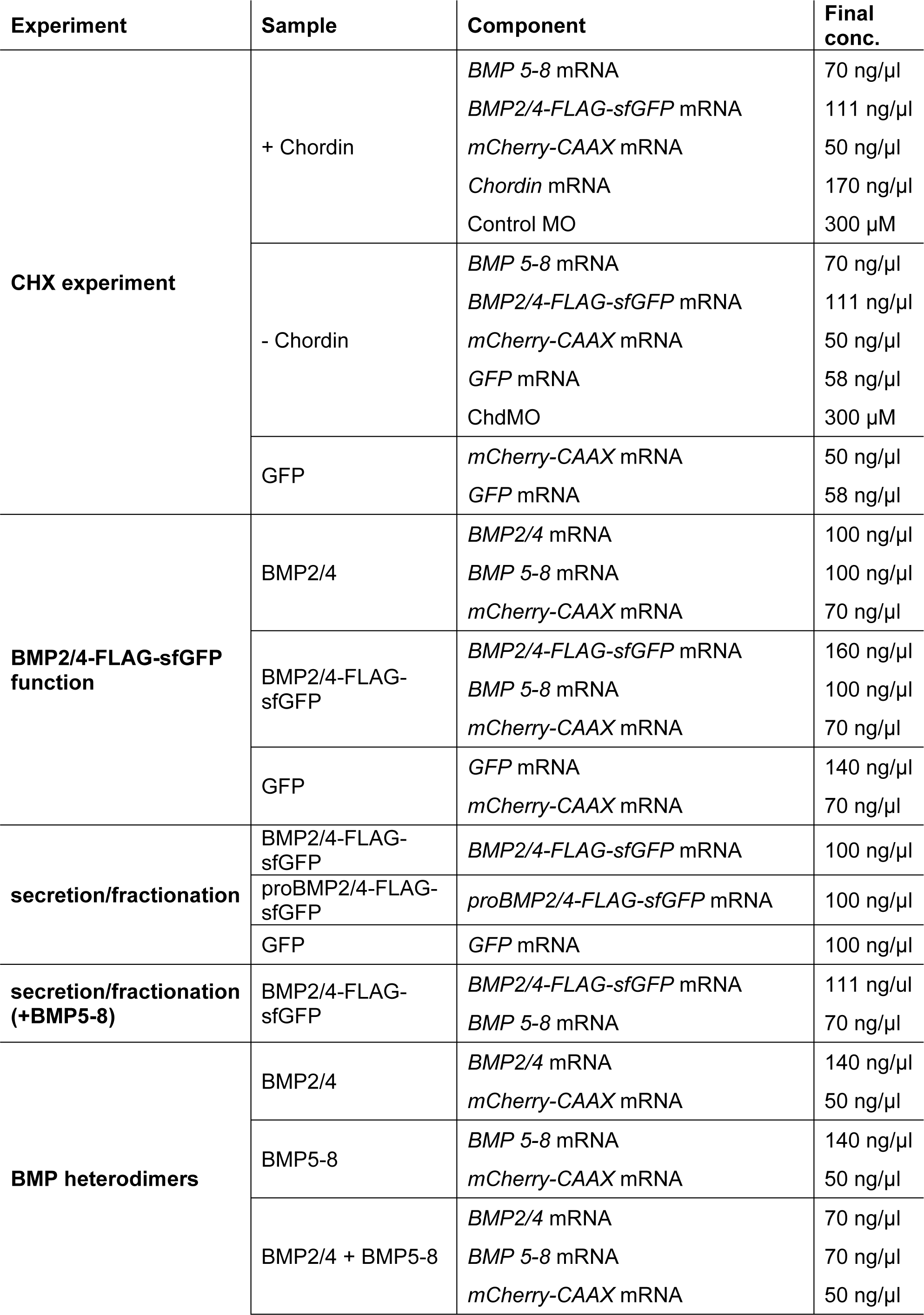
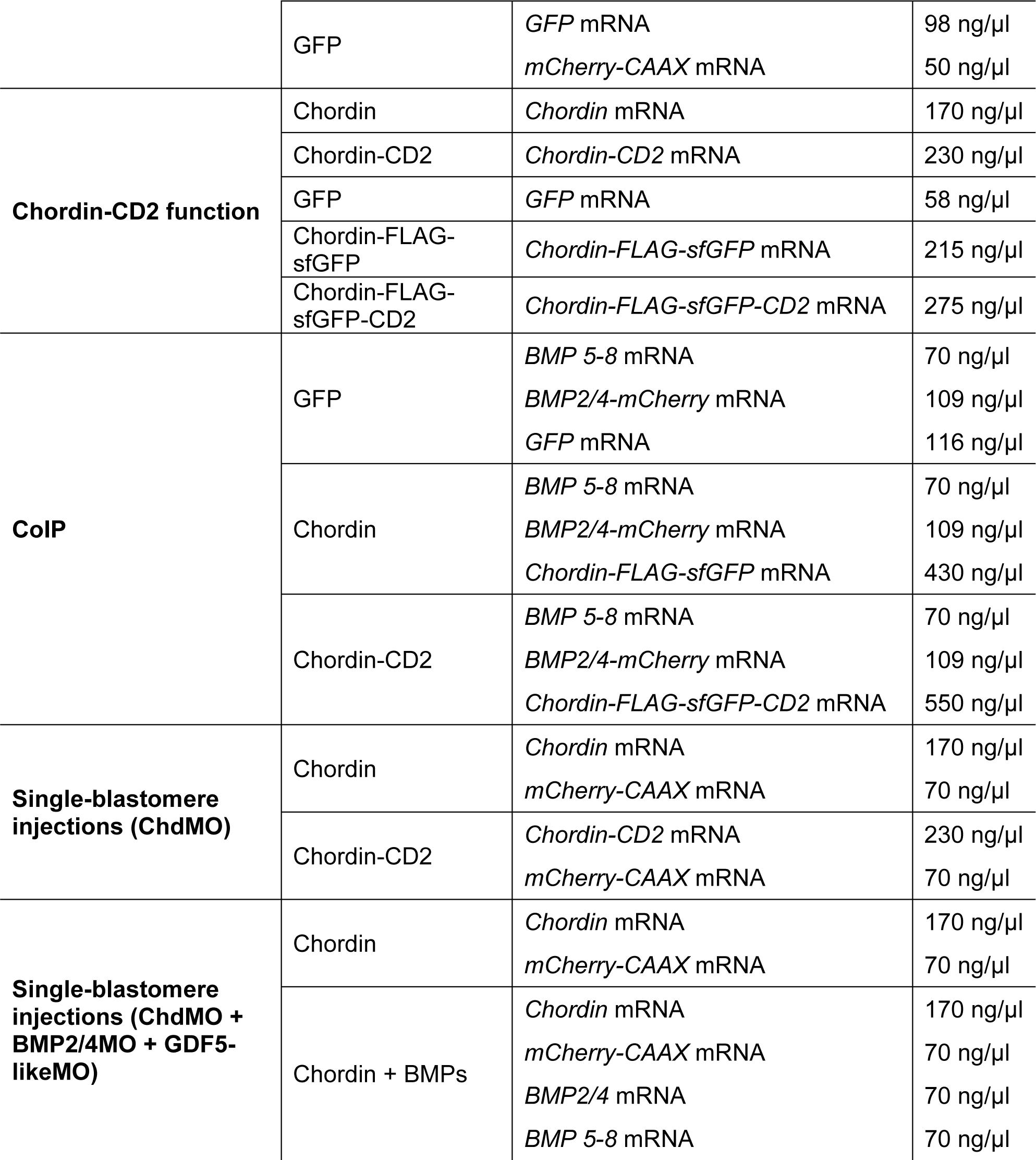
Details on concentrations used in the injection mixtures.

## Notes

### Competing Interest Statement

The authors have declared no competing interest.

